# Validation of optimised methods for avian influenza virus isolation in specific pathogen-free embryonated fowls’ eggs

**DOI:** 10.1101/2024.06.18.599543

**Authors:** Scott M. Reid, Vivien J. Coward, Joe James, Rowena D. E. Hansen, Colin Birch, Mayur Bakrania, Sharon M. Brookes, Ian H. Brown, Ashley C. Banyard

## Abstract

The internationally recognised method for diagnosis of avian influenza (AI) is virus isolation (VI) in specific pathogen-free embryonated fowls’ eggs (EFEs). In Great Britain (GB), AI virus isolation currently involves two passages in EFEs; the first typically of two days duration followed by a second lasting up to four days meaning that premises may remain under restriction for up to six days. Shorter time lengths for AIV isolation were investigated to reduce the time that businesses remain under official restrictions to safely negate AI infection, whilst maintaining test sensitivity. Both experimental inoculations of EFEs and analyses of VI attempts from high pathogenicity (HP) AI disease incursions in GB since 2016 demonstrated that HP viruses were isolated during first passage while for low pathogenicity AI outbreaks, the second passage could be reduced to two days. Power analysis showed that the benefit of reducing the number of days outweighed the risk of missing a positive isolate. This approach will substantially reduce costs to government and industry by releasing restrictions at least two days earlier where samples are negative for viral nucleic acid. Critically, it will reduce welfare implications of housing birds under restriction and improve international standards without loss of test performance.

## 1. Introduction

Avian influenza (AI) is a global threat to the poultry industry as well as to animal and human health. Safe and accurate diagnosis is a cornerstone for effective disease control. The WOAH historical ‘gold-standard’ method for diagnosis of AI is virus isolation (VI) in specific pathogen-free (SPF) embryonated fowls’ eggs (EFEs) (WOAH, 2023). Real-time reverse transcription polymerase chain reaction (rRT-PCR) in many settings has largely replaced VI but there are some applications where robust and sensitive VI is very important. The VI approach offers some advantages over rRT-PCR, namely; it demonstrates the presence of infectious virus which can be important when defining the context and relevance of results especially in investigations where all positive birds from a single case may have weak Cq values in rRT-PCR tests, that could be indicative of either early or late infection. In the latter case, assessing the data can be interpeted together with the clinical history and may fundamentally impact the decision on whether the case is proven as having actively infected birds at specific locations or premises with or without onward transmission risk. Furthermore, whilst the fidelity of rRT-PCR assays for highly sensitive and specific detection of AI virus (AIV) is proven, there is always a theoretical possibility that nucleotide subsitutions at key rRT-PCR target sites in the viral genome may compromise utility of the molecular test. Therefore, in accordance with the WOAH Manual for Diagnostic Tests and Vaccines (WOAH, 2023), VI can be used to safely exclude infection with AI in kept bird case investigations, which acts as a trigger to lift disease control measures placed upon the premises whilst diagnostic tests are conducted. These measures are designed to prevent the spread of infection until the status of the case is known.

In Great Britain (GB) our standard VI protocol requires two passages of varying duration. For notifiable avian disease (NAD; i.e., AI and Newcastle disease [ND]) investigations, quarantine/import testing and wild bird surveillance for AI, two rounds of virus propagation in EFEs are attempted, comprising an initial passage for two days (P1) followed by a second passage (P2) lasting up to four days (denoted as the ‘2 + 4’ control/standard passage model).At day two, post-inoculation into the EFE allantoic cavity, the chorio-allantoic fluid (CAF) from one of three inoculated eggs per sample is tested for haemagglutination (HA) activity. If positive by this HA test, the CAF can be further tested to determine the identity of the haemagglutinating virus but when the HA test is negative, the CAF is further passaged in EFEs. At day six total, remaining eggs are tested by the HA assay. The full duration of the current VI procedure in GB is therefore six days. During this period, the premises containing birds suspected of being infected with NAD agents remain under official disease control restrictions whilst diagnostic tests are completed.

The aim of this study was to determine whether the currently adopted two-passage model for isolation of AIV could be reduced from six days (using the ‘2 + 4’ model) without any loss of test sensitivity which would benefit GB but also provide wider international improvement. The durations were variously applied to include two, three or four days (referred to as the ‘1+1’, ‘1 + 2’, ‘1 + 3’ or ‘2 + 2’ passage models, numbers again denoting the respective P1 and P2 lengths of time (**Figure 1**)) following inoculation of AIV into EFEs. Promising outcomes from experimental studies using known virus positive original clinical material supported a reduction in the length of time from six days for completion of the protocol. In addition, we examined VI data derived from over 600 case investigations from the recent epizootics of high pathogenicity AIV (HPAIV; clade 2.3.4.4b) in GB covering the outbreak seasons 2016-2017, 2020-2021, 2021-2022 and 2022-2023, and from low pathogenicity AIV (LPAIV) cases in GB since 2014. The data was reviewed to find a correlation with the evidence base from the investigations detailed above in support of a reduction in test timing using alternative regimens.

**Figure 1.**
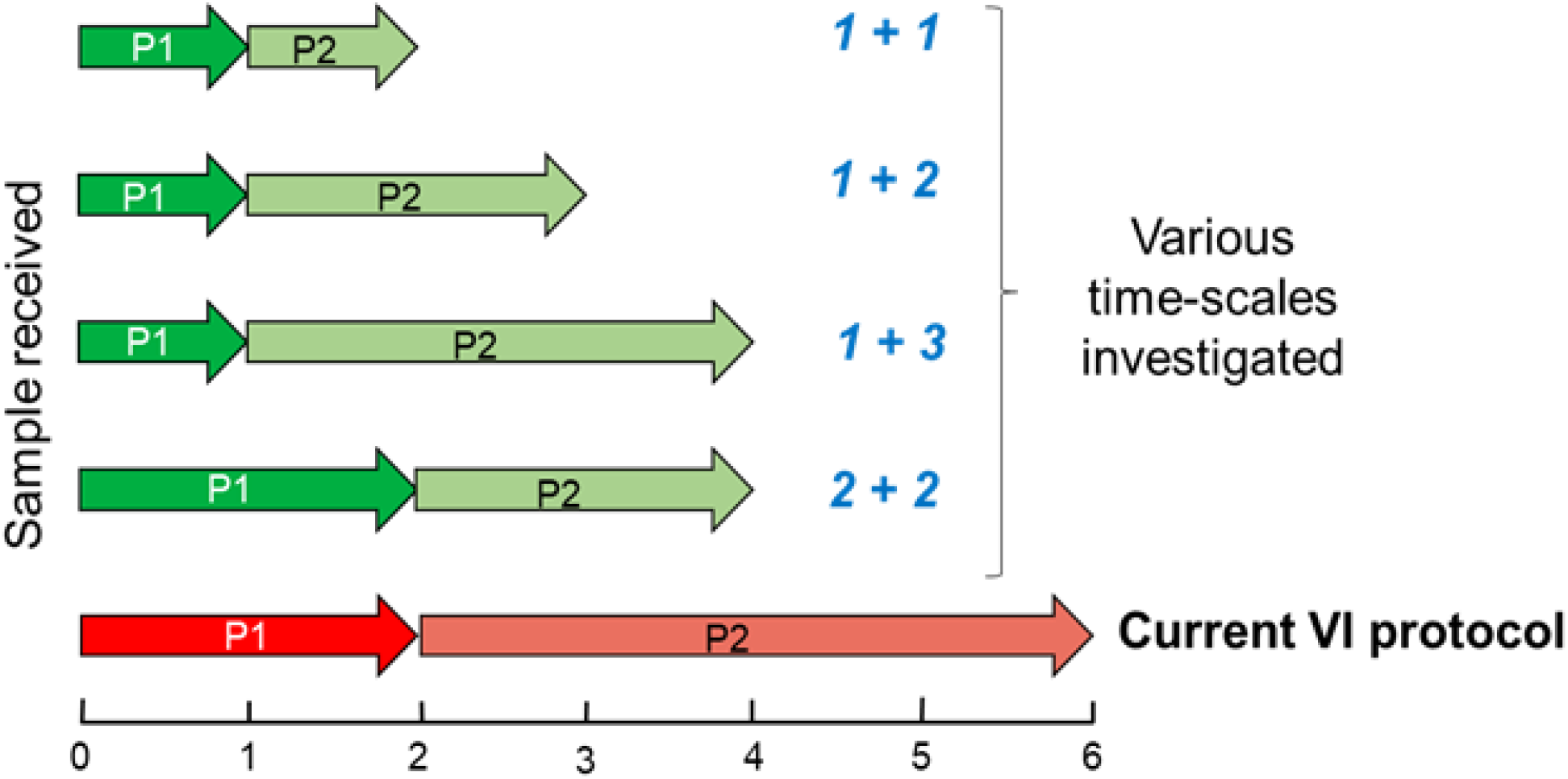
Illustration of the different approaches (‘1+1’, ‘1+2’ and ‘1+3’) for each AIV passage model under investigation, in comparison with the current (control/standard) ‘2 + 4’ model

## 2. Materials and methods

### 2.1 SPF embryonated fowls’ eggs and sample inoculation

The 9-11 days-old SPF EFEs (chicken anaemia virus-negative) used in the study were supplied from VALO BioMedia GmbH (Germany). In accordance with the legal obligations under the Animals (Scientific Procedures) Act 1986, all egg inocualtions were performed under Home Office project licences P5275AD31 (Diagnosis of Statutory and Endemic Avian Viral Diseases) and PP7633638 (Investigation of influenza virus and avian avulavirus disease), respectively in ACDP3/SAPO4 biocontainment facilities at the Animal and Plant Health Agency-Weybridge, United Kingdom.

### 2.2 Experimental inoculation of AIV from known-positive original clinical material into EFEs and determination of HA activity for each shorter passage model

Original clinical material comprising brain or intestine tissue pools from the strain HPAIV H5N8 A/chicken/England/035986/20 was inoculated undiluted into EFEs using each of the shortened models as described in **Table 1**.

**Table 1.** Daily schedule for EFE inoculations for each passage model per virus inoculum.

| Day | Minimum number of new EFEs used per model<sup>a</sup> | EFEs to undergo HA test on the day | Passage 'model' completed | Comments |
| --- | --- | --- | --- | --- |
| 0 | 5 EFEs | - | - | All passage models started |
| 1 | '1+1' - 3 EFEs<br>'1+2' - 3 EFEs<br>'1+3' - 3 EFEs | 1 | - | One EFE was passaged into 3 eggs for each model. No passage for '2+2' or '2+4' models |
| 2 | '2+2' - 3 EFEs<br>'2+4' - 3 EFEs | 4 (3 x '1+1' and 1 x day 2) | '1+1' | '2+2' and '2+4' model EFEs passage and HA test, '1+1' model EFEs undergo HA test |
| 3 | 0 | 3 (3 x '1+2') | '1+2' | '1+2' model EFEs undergo HA test |
| 4 | 0 | 6 (3 x '1+3' and 3 x '2+2') | '1+3' and '2+2' | '1+3' and '2+2' model EFEs undergo HA test |
| 5 | 0 | 0 |  | EFEs candled |
| 6 | 0 | 6 (3 x '2+4' and 3 from day 0) | '2+4' (control) | '2+4' EFEs undergo HA test |

<sup>a</sup>Total of at least 20 EFEs were required per virus inoculum for the evaluation of all passage models (including the '2+4' control/standard model).

Briefly, using the virus inoculum: Five eggs were inoculated on day 0 (P1) with 200µl of homogenate then incubated at 37^0^C overnight. Each egg batch/passage required an extra non-inoculated egg to act as a control for that specific egg batch/passage. At each passage stage, the egg embryos were decapitated as per Home Office licensure and the harvested egg fluid underwent HA testing. Shortened passage scenarios were carried out on day 1 for the ‘1+1’, ‘1+2’ and ‘1+3’ models (**Table 1**). The HA test was performed on one day 1 egg from day 0 (P1) and 200µl supernatant fluid was inoculated back into P2 (3 x eggs for the ‘1+1’, ‘1+2’ and ‘1+3’ models using a total of 9 eggs). An HA test was undertaken on the single day 2 egg (P1) and supernatant fluid inoculated back into P2 (3 x eggs for the ‘2+2’ and ‘2+4’ models; requiring a total of 6 eggs). EFEs were kept in the incubator/hot room at 37^0^C until ready for passage work and HA testing. The HA test was performed on P2 material for all models on the appropriate days according to the daily schedule (**Table 1**).

### 2.3 Isolation of virus in EFEs from samples submitted for virological investigation during HPAIV and LPAIV outbreaks occurring in GB

Assessment of samples from NAD responses was undertaken, covering the HPAIV epizootics occurring in GB during the 2016-2017 (H5N8, covering the time period from 16^th^ December 2016 to 31^st^ December 2017, **Table 2**), 2020-2021 (H5N8, 1^st^ October 2020 to 30^th^ September 2021), 2021-2022 (H5N1, 1st October 2021 to 30th September 2022) and 2022-2023 (H5N1, 1^st^ October 2022 to 30^th^ September 2023) (**Table 3**).

**Table 2.** Haemagglutinating VI data from the 13 infected premises during the 2016-2017 HPAIV H5N8 outbreak in GB.

| IP# | NAD (NDI1)<br>Reference | APHA submission reference(s)<br>for each IP | Day of isolation of<br>haemagglutinating virus<br>at P1 post inoculation<sup>a</sup> |
| --- | --- | --- | --- |
| 1 | DPR2016/31 | AV001417-16 | 1 |
| 2 | DPR2016/43 | AV000001-17 and AV000002-17 | 2<sup>b</sup> |
| 3 | DPR2017/06 | AV000039-17 | 2 |
| 4 | DPR2017/24 | AV000143-17 | 2 |
| 5 | DPR2017/35 | AV000222-17 and AV000223-17 | 2 |
| 6 | DPR2017/37 | AV000248-17 | 2 |
| 7 | DPR2017/39 | AV000281-17 | 2 |
| 8 | DPR2017/41 | AV000296-17 and AV000297-17 | 2<sup>c</sup> |
| 9 | DPR2017/51 | AV000402-17 and AV000403-17 | 2<sup>d</sup> |
| 10 | DPR2017/58 | AV000503-17 | 2 |
| 11 | DPR2017/85 | AV000851-17 | 1 |
| 12 | DPR2017/88 | AV000870-17 and AV000871-17 | 2<sup>e</sup> |
| 13 | DPR2017/101 | AV001009-17 | 2 |

<sup>a</sup> Haemagglutinating virus was isolated at P1 from the inoculated EFEs from all 13 IPs. This typically occurred on day 2 post inoculation of EFEs but virus was also isolated on day 1 as shown.
<sup>b</sup> Haemagglutinating virus isolated on day 2 from AV000001-17 but no haemagglutinating virus isolated from AV000002-17 by day 6.
<sup>c</sup> Haemagglutinating virus isolated on day 2 from AV000297-17.
<sup>d</sup> Haemagglutinating virus isolated on day 2 from both AV000402-17 and AV000403-17.
<sup>e</sup> Haemagglutinating virus isolated on day 2 from AV000870-17 but no haemagglutinating virus isolated from AV000871-17 on first passage.

**Table 3.**
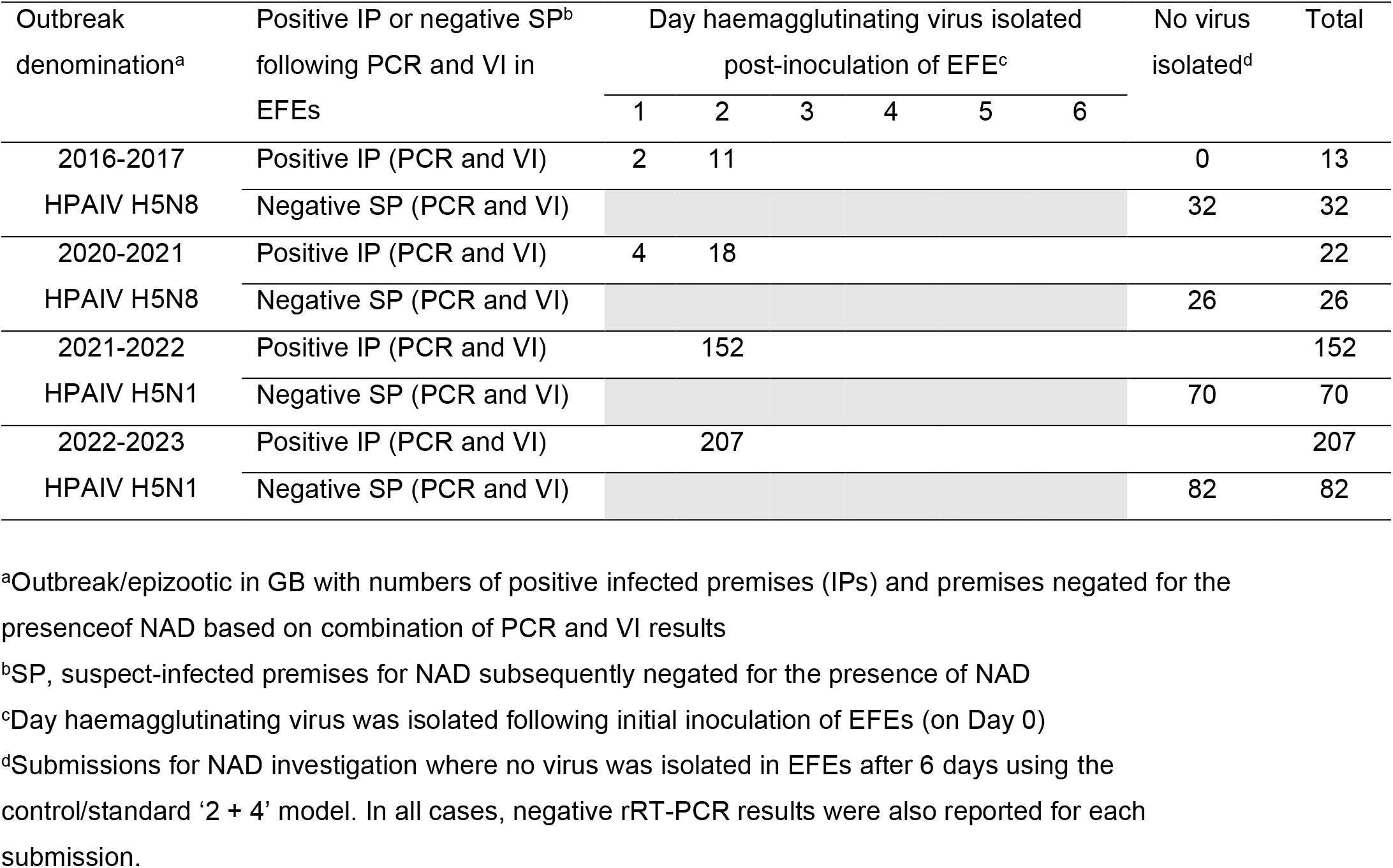
Summary of the haemagglutinating VI data from the HPAIV H5 epizootics occurring in GB since 2016.

The clinical samples from LPAIV incursions into GB since 2014, were similarly subjected to VI in 9-to 11-day-old SPF EFEs according to the internationally recognised methods (WOAH, 2023) (**Table 4**)). Typically, VI would be attempted on all sample sets comprising oropharyngeal and cloacal swabs, standard tissues (i.e., brain, lung and trachea, intestinal contents and mixed viscera homogenates) if carcasses were submitted. However, owing to the scale of the 2021-2022 and 2022-2023 AI epizootics in terms of the numbers of suspect cases from which samples were submitted for laboratory virological investigation during the peak periods of these epizootics, and the neurotropism of the H5N1 2.3.4.4b HPAIV, a streamlined algorithm was implemented and VI was often attempted only on the brain samples where carcasses were submitted in order to protect staff resource and to help to satisfy Home Office targets for reduction and refinement in the use of EFEs (NC3Rs; Russell and Burch 1959). Brain tissue was preferred in these instances as analysis of this sample type was useful in distinguishing HPAIV from LPAIV infection. For each NAD investigation, the statutory AI testing algorithm with VI, reverse transcription polymerase chain reaction (rRT-PCR) and serology described previously was employed (Reid et al., 2019).

**Table 4.** Summary of the annual VI data from the LPAIV incursions into GB since 2014.

| Incursion year<sup>a</sup> | Day haemagglutinating virus isolated post-inoculation of EFE per case<sup>b</sup> |  |  |  |  |  | No virus isolated<sup>c</sup> | Total |
| --- | --- | --- | --- | --- | --- | --- | --- | --- |
|  | 1 | 2 | 3 | 4 | 5 | 6 |  |  |
| 2014 |  | 2 |  |  |  |  | 0 | 2 |
| 2015 |  | 1 |  |  |  |  | 0 | 1 |
| 2016 |  |  | 1<sup>d</sup> |  |  |  | 0 | 1 |
| 2017 |  |  |  |  |  |  | 1<sup>e</sup> | 1 |
| 2018 |  |  |  |  |  |  | 1<sup>f</sup> | 1 |
| 2019 |  | 2 |  | 1 |  |  | 0 | 3 |
| 2020 |  | 5 |  | 1 | 2<sup>g</sup> |  | 1<sup>i</sup> | 9 |
| 2021 |  | 1 |  |  |  |  | 0 | 1 |
| 2022 |  | 1 |  |  |  |  | 0 | 1 |
| 2023 | 1 |  |  |  |  |  | 0 | 1 |

<sup>a</sup>Data shown from the respective calendar year.
<sup>b</sup>Day haemagglutinating virus was isolated following initial inoculation of EFEs (on Day 0). All viruses isolated by day 2 post-inoculation of EFEs were from P1 eggs.
<sup>c</sup>Submissions for NAD investigation where no virus was isolated in EFEs after 6 days using the control/standard '2 + 4' model.
<sup>d</sup>Haemagglutinating virus was isolated on P2.
<sup>e</sup>LPAIV H5N3 detected by molecular methods only.
<sup>f</sup>LPAIV H6N5 detected by molecular methods only.
<sup>g</sup>Haemagglutinating virus harvested on day 5 but eggs not checked on days 3 or 4.

For each suspect HPAI or LPAI case investigation, the VI methodology essentially followed the control/standard ‘2 + 4’ model whereby official clinical samples were inoculated into each of three eggs for incubation at 37°C (P1). At day two, post-inoculation into the EFE allantoic cavity, the CAF from one of the three eggs per sample was tested for HA activity. When the HA test was negative, the CAF was further passaged into three EFEs (P2). At day six total, all eggs were tested by the HA assay (**Figure 1**). However, in line with Home Office project licence legislation, all EFEs were candled on day 1 post-inoculation. From 2022, if the rRT-PCR results were positive for the submission, one egg was typically taken off for HA testing per submission on day 1. Otherwise, on day 2 post-inoculation, one egg was taken off per sample and subjected to HA testing. In the event of haemagglutinating virus being detected on either day 1 or day 2 (P1), at least one isolate was harvested, placed into storage and the VI test concluded by chilling all the remaining eggs inoculated for the submission at 4°C according to the Home Office regulations.

In the absence of haemagglutinating virus following HA testing for the submission on day 2, the egg harvest from each sample (P1) was inoculated back into three fresh EFEs for incubation for up to four more days at 37°C (P2) along with the remaining one or two EFEs for each sample (**Figure 1**). All EFEs were subsequently candled on day 3 as this corresponded to day 1 post-inoculation of P2 eggs. Eggs were also candled on day 4 post-inoculaton if the P1 eggs reached 14 days of age. HA testing was then performed on those eggs showing signs of embryo death to determine whether this was due to haemagglutinating virus. Furthermore, in strict adherence to the Home Office licence regulations, EFEs had to be candled daily from 14 days of age to check for egg death.Therefore, in the absence of haemagglutinating virus in the EFEs up to that time point, all inoculated EFEs remaining alive had to be candled on day 5 and subjected to HA testing on day 6 (**Figure 1** and **Tables 2-4**).

## 3. Results

### 3.1 Virus isolation in EFEs using known-positive original clinical material with variable passage parameters

Following EFE inoculation with the dilution series of HPAIV H5N8 A/chicken/England/035986/20, with the ‘2+2’ and ‘2+4’ (control/standard) models, re-isolation of virus at P1 was successful, eliminating the need for an additional passage. Positive haemagglutinating virus was recovered with the reduced ‘1+2’, ‘1+3’ and ‘2+2’ passage models but not for the ‘1+1’ model. As all P1 eggs had died at day 2, no P2 egg harvests were set up. All P2 eggs set up on day 1 were dead with the ‘1+1’ or ‘1+2’ models (data not shown).

### 3.2 Statutory isolation of HPAIV and LPAIV in EFEs during AI outbreak events in GB

#### 3.2.1 2016-2017 HPAIV H5N8 (clade 2.3.4.6) epizootic in GB

**Table 2** shows the time periods (number of days) post inoculation into EFEs for isolation of HPAIV H5N8 from each of the 13 IPs comprising the 2016-2017 HPAIV H5N8 outbreak in GB. The time period of this epizootic was defined from 16^th^ December 2016 up to 31^st^ December 2017. In all cases, haemagglutinating virus was isolated at P1 (without passage) by day 2 post-inoculation of EFEs. Second passage and later timepoints were not investigated as the eggs were already positive for each submission.

#### 3.2.2 2020-2021 (H5N8), 2021-2022 (H5N1) and 2022-2023 (H5N1) HPAIV epizootic seasons

**Table 3** further summarises the VI data from the 13 IPs from the 2016-2017 HPAIV H5N8 outbreak along with the summary of the time periods (days) for isolation of haemagglutinating virus post inoculation into EFEs from samples submitted from the IPs over the 2020-2021 (H5N8), 2021-2022 (H5N1) and 2022-2023 (H5N1) HPAIV epizootic seasons.

In all cases where haemagglutinating HPAIV was successful (n=394), an isolate was harvested at the P1 stage either on day 1 or day 2 post-inoculation of the EFEs. The positive VI result from all IPs was also supported by the positive results from the statutory rRT-PCR testing of the official clinical samples submitted from the respective IP and from the gross pathological examination when carcasses were submitted for investigation along with swabs (data not shown).

#### 3.3.3 LPAIV incursions into GB from 2014 to 2023

The annual VI data from the IPs associated with the LPAIV incursions into GB from 2014 to 2023 is summarised in **Table 4**. LPAIV incursions included non-H5 and non-H7 AIV cases.

Out of 16 VI-positive cases stemming from the LPAIV incursions, haemagglutinating virus was detected by day 2 (on P1) in all but three cases (**Table 4**). With two cases (both occurring in 2020), harvesting of virus occurred on day 5 as the inoculated eggs were not examined for haemagglutinating activity on days 3 and 4. The data from both cases was therefore not included in the analyses. Virus isolates were obtained on either day 3 or day 4 post-inoculation on three occasions which supported our hypothesis that the second passage can be reduced from four days to two days. Overall, the data further supports the use of shortened passage models covering up to 4 days in duration with passage.

## Discussion

Whilst PCR has largely replaced VI in many applications where definitive detection of AIVs is required, this method nevertheless is still used as an international ‘gold’ standard. There are circumstances where confirmation or exclusion of the presence of infectious AIV is important to a case investigation. One such application is the ability to reliably negate suspicions of disease due to AIVs in kept birds that have already returned preliminary PCR results indicating absence of virus. Virus isolation has some advantages over PCR when seeking to confirm active infection but the time to obtain such results is laborious and time consuming using methodology that was developed many decades ago. Therefore, in this study we aimed to review the methodology and investigate if alternative designs (models) could achieve faster results but with the same level of test sensitivity. Consequently, a science evidence base has been generated to support a revision to the internationally defined diagnostic methods for isolation of AIV prescribed by WOAH (WOAH, 2023). We report that a negative VI result for AIV in EFEs can be confirmed by day 4 (with passage) following inoculation of clinical samples into EFEs, rather than six days as is currently prescribed using the control/standard ‘2 days + 4 days’ model.

Analysis of the VI passage data from statutory NAD investigations during the epizootics of HPAIV H5N1 and H5N8 (clade 2.3.4.4b) occurring in GB since 2016 showed that where VI was successful, the isolate was generated from the P1 EFEs (day 1 or day 2 post-inoculation) (**Table 2** and **Table 3**). **Table 3** shows that all 394 of the HPAIV-positive isolates generated were detected within day 1 or 2 of the first passage. There were 210 examples of ‘no virus isolations’ in either the first passage (days 1-2) or the second passage (days 3-6). In **Table 4**, which contains isolation data for LPAIV cases, 13 positive isolations were made in the first passage and 3 more isolations were made in the first two days of the second passage. The two isolations on day 5 were discarded as these eggs were not checked on day 3 or day 4. There were 3 examples of ‘no virus isolations’ after the 6 days. By considering only the positive isolations, a retrospective power analysis was carried out to determine the probability that a positive sample would be isolated in day 1-4 (the 2+2 model) rather than in day 5 or 6 (the 2+4 control/standard model). By using the 410 cases of positive isolations that occurred in day 1-4 compared to 0 positive isolations that occurred on day 5-6, and a 95% confidence interval, we calculated this probability as 98%. This analysis showed that the benefit of reducing the number of days outweighed the risk of missing a positive isolate.

Experimental inoculation of EFEs with known positive original clinical material provided further evidence to support the adoption of the shortened passage model from six to four days duration with passage. Importantly, the viruses from statutory NAD investigation activities in the field (i.e., original clinical samples) were not necessarily well adapted to eggs as these came from multiple different avian species, and so may not efficiently and productively replicate in EFEs. Hence, the observation that all fresh clinical samples from naturally infected birds that contained virus (determined from rRT-PCR testing), submitted for investigation from across multiple outbreak seasons in GB were viable with respect to VI within the four day timeframe, thereby supporting this proposed reduction in the passage protocol (**Table 3** and **Table 4**). Furthermore, the infectivity of material inoculated from field samples may vary according to multiple factors such as the condition of the birds when samples were taken and temperature during transport which could influence the likely rate of initial rapid virus replication from the samples. Nevertheless, despite these obstacles to successful VI, the outbreak data fully supports a reduction in protocol length to 4 days with passage without the loss of test sensitivity.

The optimisation of this methodology provides an evidence base to modify international standard approaches for VI of AIV. If international agreement is achieved this will have a profound impact upon the kept bird industry in instances where premises are negative using molecular tools but international standards require VI to be completed. While it is extremely difficult to align costs against the benefit of reduced period assigned to VI testing for AIV, the impacts are likely to be substantial for both bird and human welfare. From an avian perspective, a reduction in time where disease control restrictions are in place will impact upon flocks that are negative for NAD. Welfare implications in these situations can be considerable, especially where flocks reach age and size capacity within housing. Furthermore, being under disease control restrictions can have a knock on effect with downstream processes. For example, a reduction in the positive/negative virus detection reporting time length also benefits chicks in hatcheries with no ‘home’ to go to, as the sites of destination may still contain birds, or have not been able to carry out effective cleansing and disinfection ready to receive new birds. The financial cost and emotional burden felt by owners of poultry in this situation are considerable. Costs incurred when the affected premises are placed under disease control restricton include losses from untreated, sick birds; losses due to birds not being able to be processed appropriately according to age and size, and losses from birds not being available for the correct product/season including birds growing too large to fulfil order requirements and birds missing key market dates, for example. Therefore, despite the unavailability of cost benefit financial data, these factors mean that the costs are already significant on any business as well as the emotional stresses placed on farm staff. The evidence gathered in this study substantially impacts upon all features of the sector and will potentially have global outreach if it is agreed it provides a robust evidence base to safely change method design. Data regarding negation for ND is being gathered in parallel to further support a change in this methodology that encompasses all NADs, since by definition disease suspicions cannot safely exclude one disease from the other and both are relevant for exclusion of exotic avian viral disease.

The successful outcomes from these analyses collectively supported a reduction in the length of time businesses/premises remain under official restrictions when investigating suspicions of AI, thereby substantially reducing costs both to the government and industry through earlier detection or negation of infection, with concomitant reduction in impact on animal welfare with the shortened VI test turnaround times. In addition to the financial savings, of particular significance in reducing the total time length of the shorter passage model is the potential to lift disease control restrictions on the affected premises following a negative (i.e., ‘no virus isolated’ from a negative HA test result) two-day passage result at P1 following inoculation of the original clinical samples, or on collected allantoic fluids at EFE passage in tandem with negative rRT-PCR results.

## Acknowledgements

Financial support was provided by Defra, and the devolved administrations of Scotland and Wales through contracts SV3400 (Monitoring for statutory and exotic virus diseases of avian species) and SE2213 (FluFutures 2.0 - Understanding the diverse spectrum influenza virus based threats to the UK). The authors also acknowledge Dr Zoe Treharne, Veterinary Lead of the Avian Species Expert Group at APHA and Maire Burnett of the British Poultry Council for their assistance.

## Ethical issues

The described VI using SPF EFEs was performed according to the internationally recognised method described by WOAH under Home Office Project Licences P5275AD31 (Diagnosis of Statutory and Endemic Avian Viral Diseases) and PP7633638 (Investigation of influenza virus and avian avulavirus disease).

## References

WOAH (World Organisation for Animal Health), 2023: Manual of Diagnostic Tests and Vaccines for Terrestrial Animals;

Chapter 3.3.4. “Avian Influenza (including infection with high pathogenicity avian influenza viruses)”, (Version adopted in May 2021). Available at: https://www.woah.org/fileadmin/Home/eng/Health_standards/tahm/3.03.04_AI.pdf

Russell WMS, Burch RL, 1959 (as reprinted 1992): The principles of humane experimental technique. Wheathampstead (UK): Universities Federation for Animal Welfare.

Reid SM, Núñez A, Seekings AH, et al. Two Single Incursions of H7N7 and H5N1 Low Pathogenicity Avian Influenza in U.K. Broiler Breeders During 2015 and 2016. Avian Dis. 2019 Mar 1;63(sp1):181–192. doi: 10.1637/11898-051418-Reg.1. PMID: 31131576.

